# A bacterial parasite effector mediates insect vector attraction in host plants independently of developmental changes

**DOI:** 10.1101/036186

**Authors:** Zigmunds Orlovskis, Saskia A Hogenhout

## Abstract

Parasites can take over their hosts and trigger dramatic changes in host appearance and behaviour that are typically interpreted as extended phenotypes to promote parasite survival and fitness^1^. For example, *Toxoplasma gondii* manipulates the behaviour of infected rodents to aid transmission to cats^2^ and parasitic trematodes of the genus *Ribeiroia* alter limb development in their amphibian hosts to facilitate predation by birds^3^. Plant parasites and pathogens also reprogram host development and morphology^4^. Phytoplasma parasites of plants induce extensive leaf-like flower phenotype (phyllody) in their host plants, presumably to attract insect vectors on which these bacteria depend for transmission^5,6^. However, it remains debatable whether morphological phenotypes, such as phyllody, are directly beneficial to the parasites or are side-products of parasite infection^7,8^. Previously, we found that phytoplasma virulence protein (effector) SAP54 binds and mediates degradation of host MADS-box transcription factors 26 (MTFs), regulatory hubs of plant development and hormone physiology, to induce phyllody and promote insect vector colonisation^5^. Here we show that plants heterologously expressing SAP54 are strongly attractive to insects, but surprisingly, insect attraction was independent of the presence of leaf-like flowers. Moreover, plants that produce leaf-like flowers in the absence of SAP54 did not attract insects. We conclude that the SAP54 effector mediates insect vector attraction in host plants by exploiting the role of its MTF targets in insect defence and that perturbation of floral development may be a secondary effect of the effector activity.

## Results and Discussion

The aster leafhopper *Macrosteles quadrilineatus* is the most important insect vector of the phyllody-inducing ‘Ca. Phytoplasma asteris’ strain Aster Yellows Witches Broom (AY-WB). This leafhopper favours to colonise phytoplasma-infected plants and GFP-SAP54 transgenic plants with phyllody/leaf-like flowers compared to non-infected plants and GFP transgenic plants with wild type flowers ^5^ We confirmed these findings in independent insect choice experiments; the insects produced more progeny on GFP-SAP54 transgenic plants with leaf-like flowers (Figure 1A) and, in addition, we found that the insects also spent more time on plants with leaf-like flowers (Figure S1), thus demonstrating both reproductive and orientation preference of insect vectors for these plants. However, when insects were not given a choice between host plants, by caging the leafhoppers on either GFP-SAP54 transgenic plants with leaf-like flowers or control GFP transgenic plants with wild type flowers, no increase in nymph production was observed (Figure S2).

**Figure 1.**
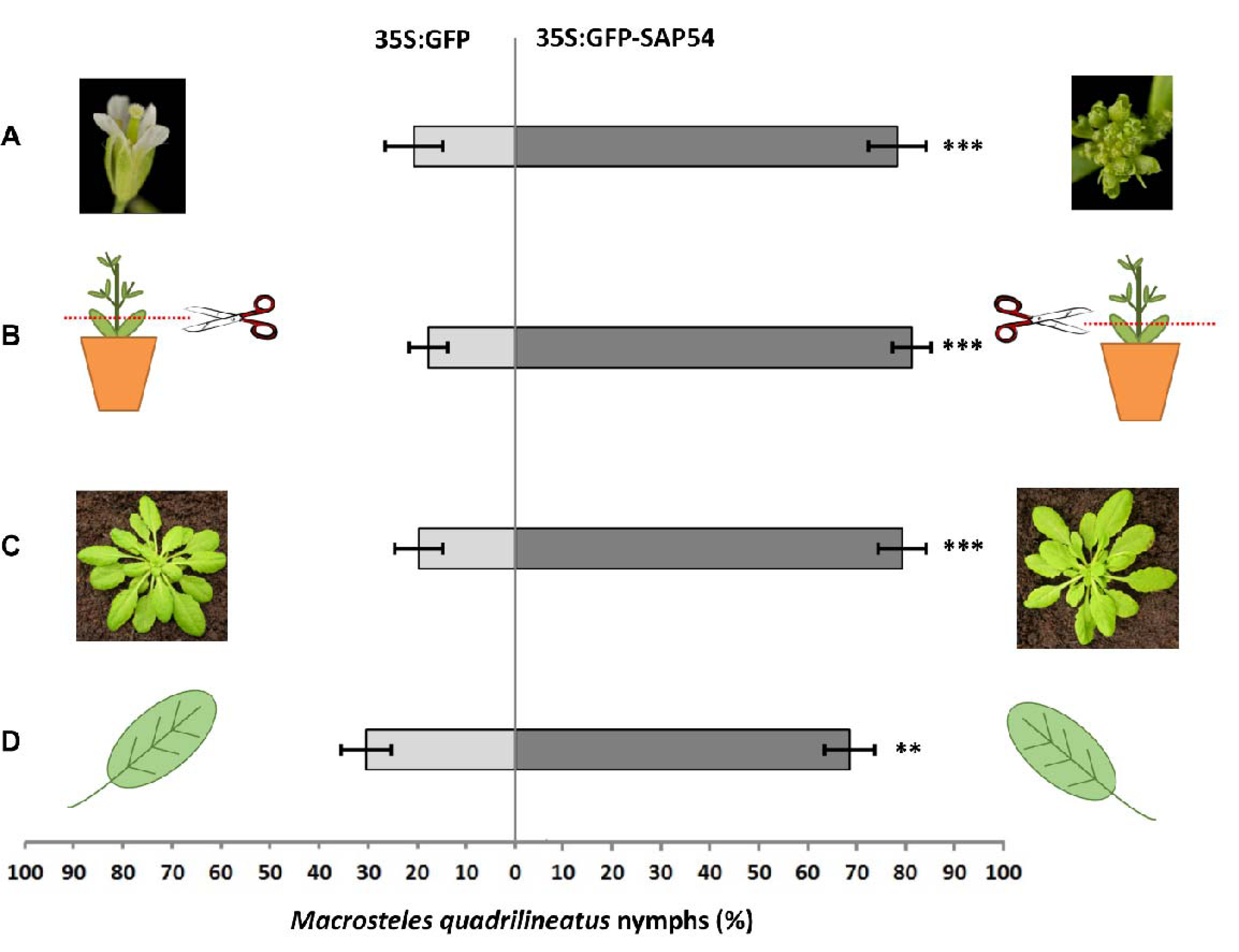
Flowers and transition from floral to vegetative phase are not required for SAP54-mediated enhancement of insect colonization. **(A)** *M.quadrilineatus* produces more nymphs on 35S:GFP-SAP54 transgenic *A.thaliana*(Col-0) plants with leaf-like flowers than on 35S:GFP (Col-0) control plants with wild type flowers (p<0.001). **(B)** Removal of flowers and floral stems does not affect *M.quadrilineatus* colonization preference of 35S:GFP-SAP54 transgenic *A.thaliana*(Col-0). **(C)** Leafhoppers also prefer GFP-SAP54 transgenic plants prior to transition from vegetative to floral growth (plants grown under short day photoperiod 10h/14h light/dark). **(D)** *M. quadrilineatus* lays more eggs on single rosette leaves of GFP-SAP54 transgenic plants. Except for C, all other choice tests were conducted with 6-week old plants grown at 22^o^C, 16/8-hour light/dark. For A-C, 10 male and female leafhoppers were offered to choose between opposite-facing whole plants for 5 days. Leafhoppers were removed and the individual plants were bagged. The total number of nymphs per plants in each test cage were counted two weeks later. In D, leafhoppers were allowed to chose between equally sized rosette leaves from opposite-facing 35S:GFP and 35S:GFP-SAP54 plants for 5 days. Upon removal of the leafhoppers, eggs laid on the leaf surface were counted. Bars show the mean percentages ±SEM of nymphs/eggs per total number of nymphs/eggs per test cage at n = 6 cages. *** p<0.001; ** p<0.025 (paired t-test). All experiments were repeated three times with similar results.

Thus, the observed leafhoppers preference is the result of preferential orientation to plants with leaf-like flowers rather than an increase in reproductive efficiency *per se.*

These results prompted us to further examine if the insects are attracted by leaf-like flowers or repelled by wild-type flowers. Interestingly, insects resided and occasionally fed on both normally developed as well as leaf-like floral structures (Figure S3A), suggesting that the two types of flowers do not attract neither repel the insects. In addition, we noticed, that most of the insects preferred to reside on the rosette leaves rather than floral stems or flowers (Figure S3B), suggesting that the flowers are not required for leafhopper attraction. To analyse the impact of leaf-like flowers on leafhopper preference further, we removed both the leaf-like and wild-type flowers from plants in the insect choice experiments and found that the leafhoppers also preferred the GFP-SAP54 plants without leaf-like flowers (Figure 1B). *A. thaliana* plants used in insect choice tests so far were grown at long days to induce bolting and flowering. Next, we conducted choice tests on *A. thaliana* plants grown at short days and that had vegetative organs (rosettes) without flowers. Again, *M. quadrilineatus* produced more nymphs on GFP-SAP54 versus GFP (control) plants at the vegetative growth stage (Figure 1C), suggesting that not only changes in floral morphology but also physiological and developmental transformations during floral transition are not necessary for insect attraction. To confirm this finding, leafhoppers were also given a choice between single leaves of GFP-SAP54 and GFP plants. We found that the leafhoppers preferred to lay eggs onto single leaves of GFP-SAP54 plants (Figure 1D), indicating that leafhoppers are attracted to the leaves. Taken together, these data suggest that leaf-like flowers are not required for host plant selection by the leafhopper vector, and that the SAP54-mediated modulation of plant vegetative tissues rather than plant reproductive organs is involved in insect vector attraction.

The above experiments provide evidence that leaf-like flowers are not required for insect vector preference. Nonetheless, these flowers could contribute to the insect preference. To test this, we conducted choice experiments with *A. thaliana* lines displaying leaf-like flowers, including MTF mutant lines *ap1*^9^ and *lfy*^10^ and the 35S:SVP transgenic line ^11^. All these lines produce flowers that share leaf-like structures reminiscent to those of phytoplasma-infected and GFP-SAP54 transgenic plants ^6^. We found that leafhoppers produce similar numbers of progeny on both plants indicating no colonization preference for either wild type or mutant plants with leaf-like floral phenotypes (Figure 2). These data are in agreement with insect preference for rosette leaves rather than floral stems or flowers (Figure S3B). Thus, the leaf-like flowers are neither required nor involved in the leafhopper vector preference.

**Figure 2.**
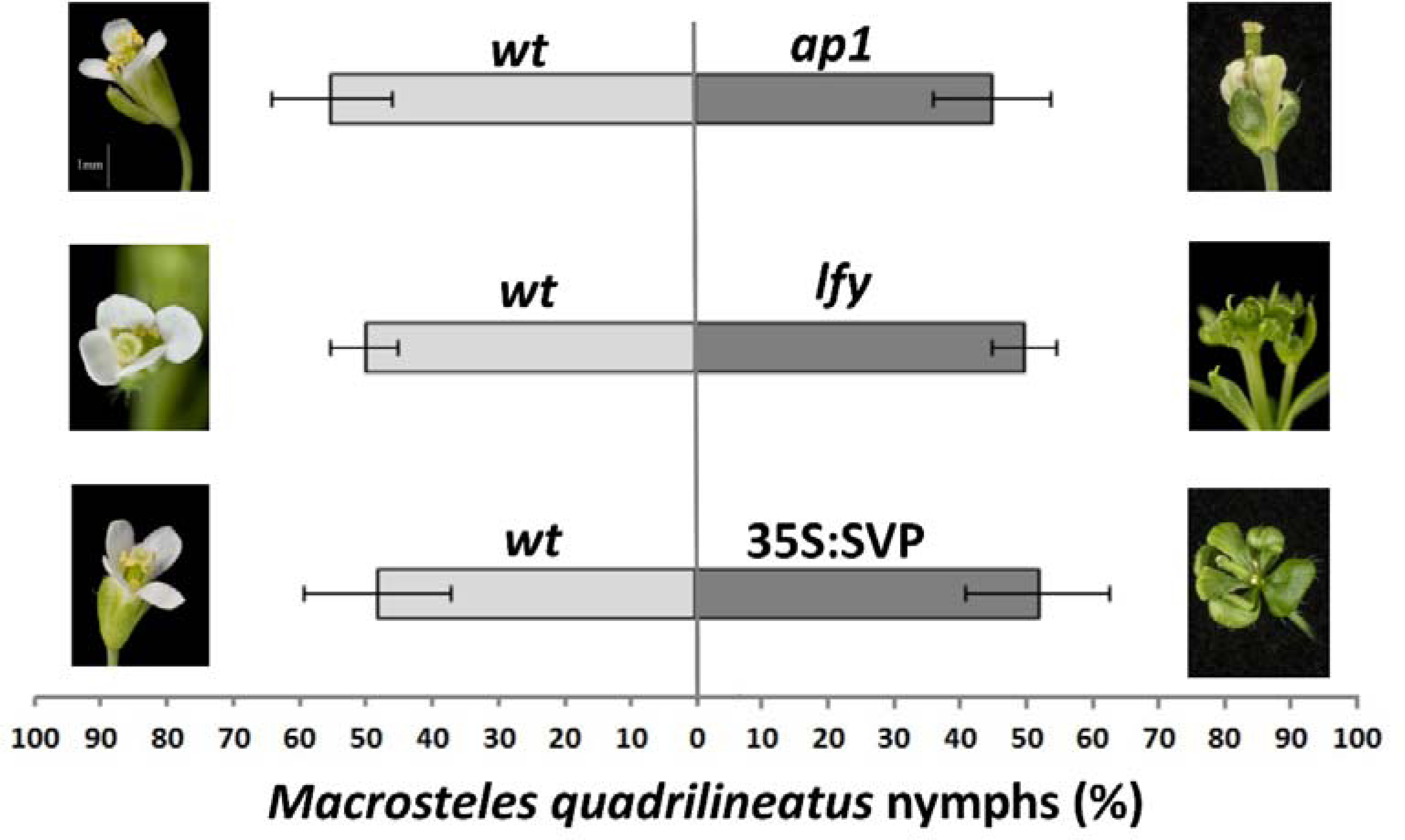
Aster leafhopper *Macrosteles quadrilineatus* has similar oviposition preference for plants with normal and leaf-like flower phenotype. *M. quadrilineatus* did not show a preference for colonization of Col-0 wild type versus Col-0 *apetala1 (ap1-12)* (p = 0.835), Col-0 versus Col-0 *leafy (lfy-1)* (p=0.985) and Col-0 versus 35S:SVP (Col-0) (p=0.960). Experiments were conducted with whole plants as described in the legend of Fig. 1. Bars shows percentages ±SE of living *M.quadrilineatus* nymphs found on each test plant per total number of nymphs within a single choice cage. Data were analysed by paired t-tests. All experiments were repeated three times with similar results.

Hitherto, direct analyses of the adaptive significance of parasite extended phenotypes have been limited because many parasites (such as phytoplasma) are not amenable to genetic manipulation and parasite genetic factors that induce the dramatic host alterations are often unknown. Here we used mechanistic knowledge of the phytoplasma virulence protein SAP54 to dissect if phytoplasma-induced phenotypic changes of plant hosts, including phyllody and insect vector attraction, are connected. We previously demonstrated that SAP54 interaction with the plant 26S proteasome cargo protein RAD23 is required for both the induction of leaf-like flowers and insect vector attraction ^5^ supporting the hypothesis that phytoplasma-induced morphological changes in plants such as leaf-like flowers may be required for insect attraction ^12,13^. However, this study has shown that leaf-like flowers are not required nor are involved in insect vector attraction. Moreover, leafhoppers preferred plant vegetative tissues over reproductive organs. Thus, leaf-like flowers do not promote leafhopper colonization, even though these two phenotypes are genetically connected via SAP54 interaction with the 26S proteasome cargo protein RAD23 ^5^.

Phyllody-inducing ‘*Ca.* Phytoplasma asteris’ phytoplasmas, such as AY-WB, often infect annual plants that die upon completion of their life cycle ^14^ Phytoplasmas are dependent on insect vectors for leaving the plant before it dies ^13^ Thus, insect attraction promotes fitness of phytoplasma and therefore it is likely that insect vector attraction is the extended phenotype of SAP54. A role of leaf-like flowers in phytoplasma fitness has become less clear. It is possible that the induction of phyllody is a side effect of SAP54-mediated modulation of processes involved in insect attraction. SAP54 induces leaf-like flowers by mediating degradation of MTFs via interaction with RAD23 ^5^. MTFs are regulatory hubs for a plethora of physiological processes in plants, including plant immunity (comparable to animal HOX genes); several MTFs appear to (in)directly regulate cytokinin and jasmonic acid (JA) synthesis and response genes ^11^, which affect plant-insect interactions ^15-18^, such as that of the AY-WB leafhopper vector *M. quadrilineatus*^19^. In addition, MTFs regulate age-related resistance responses to pests ^20^. Therefore, SAP54-mediated degradation of MTFs may modulate plant immunity leading to attraction of the leafhoppers.

Another AY-WB phytoplasma effector, SAP11, binds and destabilizes specific members of the TEOSINTE BRANCHED1, CYCLOIDEA, PROLIFERATING CELL FACTORS 1 and 2 (TCP) family and promotes leafhopper oviposition activity in no-choice tests ^19,21,22^. TCPs are transcription factors conserved among plants that are regulatory hubs for plant growth and organ formation, and in addition, regulate a variety of microRNAs and the plant defence hormones jasmonic acid (JA) ^23,24^ and salicylic acid (SA) ^25^. Another phytoplasma effector, TENGU, also induces witch's broom-like symptoms in plants ^12^ and alters the plant JA and auxin hormone balance ^26^, and it was suggested that the witch's broom-like symptoms attract the leafhopper vectors ^12^. However, given that SAP11 decreases JA production, which increases leafhopper colonization ^27-29^, it remains to be investigated if witch's broom symptoms are involved in the leafhopper colonization preference. Thus, the SAP54, SAP11 and TENGU effectors all alter plant development that resemble symptoms of phytoplasma-infected plants, but for SAP54 we have now shown that the alterations in plant development (leaf-like flowers) are not required for insect preference.

Targeting conserved plant proteins, such as MTFs and TCPs by phytoplasma effector proteins may enable the phytoplasma parasites to infect a broad range of plant species. The 26S proteasome shuttle proteins RAD23 are also conserved among plant species ^30^ Compatibility of phytoplasmas with multiple plant species is essential given that AY-WB phytoplasma and related parasites are transmitted by polyphagous insect species of the genus *Macrosteles*^13,14^. Because these insects readily feed on many plant species, phytoplasmas will increase their fitness if they can modulate these plants to increase attraction and colonization of insect vectors. In agreement with this, SAP54 homologs are found in diverse phyllody-inducing phytoplasmas that infect a wide range of plant species ^6,31,32^. Thus, generalist parasites, especially those dependent on alternative hosts for transmission, could gain fitness benefits via interfering with conserved host processes.

The adaptationist view is that parasite-induced changes in hosts are selected for the benefit of the parasite ^1,33^. Alternatively, the activity of parasite genes may also lead to the emergence of non-adaptive secondary phenomena ^7^. In agreement with the latter, phyllody in phytoplasma-infected plants may have been selected together with the primary adaptive role of SAP54 in enhancing insect vector attraction. Like phytoplasmas, other parasites induce a complex package of changes in hosts, some of which are viewed as adaptive or neutral with respect to selective pressures. For example, alterations in limb morphology and number in trematode *(Ribeiroia* species) infected frogs may or may not be the primary evolutionary mechanism for vertical transmission to birds ^3,34^. Similarly, the adaptive significance of the leaf morphological changes induced by gall-forming insects remains to be tested and is subject to various alternative hypothesis ^35,36^, although the adaptive explanations tend to be preferred. The non-adaptive explanation and the adaptationist view are equally instrumental in understanding the evolution of parasite-altered host phenotypes and mechanistic insights into the functions of parasite virulence genes, as we did for SAP54, are key to uncouple the two.

## Experimental Procedures

### Generation of plants for insect assays

Generation of 35S:GFP-SAP54 and 35S:GFP transgenic *Arabidopsis* lines was done according to methods described in ^6^. *A. thaliana ap1* and *lfy* mutant were obtained from NASC (ID: N6232, allele ap1-12; ID: N6228, allele lfy-1). The 35S:SVP lines were kindly provided by Martin Kater and described in ^11^. The rad23 mutant lines were provided by Richard Vierstra and described in ^30^. To generate infected plants, five-week old plants were infected with ‘*Ca.* Phytoplasma asteris’ strain Aster Yellows Witches Broom (AY-WB) by adding five AY-WB-carrier adult *Macrosteles quadrilineatus* Forbes (Hemiptera: Cicadellidae) to each plant in a transparent Perspex tube (10cm high, diameter 4cm) for inoculation access period of 5 days. Three rosette leaves were collected for extraction of genomic DNA to confirm phytoplasma infection using AY-WB specific primers BF 5’ AGGATGGAACCCTTCAATGTC 3’ and BR 5’ GGAAGTCGCCTACAAAAATCC 3’ ^5^. All plants used in insect choice experiments were sown on insecticide-free F2 compost soil (Levington). In order to stimulate flowering, the test and control plants were transplanted in 10cm x 10cm square pots (F2 soil) and grown for 3 weeks at 22°C, long day photoperiod (16/8-hour light/dark). For the experiments involving no-flower formation, plants were grown at 22°C in short day photoperiod (10/14-hour light/dark) for 8 weeks.

### Insect choice assay

All insect choice experiments were performed in transparent polycarbonate cages 62cm x 30cm x 41cm (H x W x D). Two opposite sides of the cage were fitted with white nylon mesh held by magnetic strips to the carcass of the cage for ventilation and access. Two test plants infected with AY-WB 21 days earlier were placed randomly diagonally opposite each other in the corners of a cage. 10 male and 10 female adult *M. quadrilineatus,* which did not carry AY-WB phytoplasma, were released from a transparent Perspex tube (9cm high, diameter 3cm) in the centre of the cage, at equal distance from each test plant. Adult insects were removed 5 days after addition to the cage. Plants were removed from the choice cage and contained individually in transparent perforated plastic bags at 22°C, long day photoperiod (16/8-hour light/dark). Nymphs were counted on each test plant 14 days after removal of adult insects from the cages. Data were expressed as proportion of total number of nymphs found on the test plants within each choice cage.

### Insect no-choice assay

For no-choice experiments 5 female and 5 male non-infected adult *M. quadrilineatus* were added to individual plants surrounded by a transparent plastic cage. Plants were grown and insect progeny measured as in choice experiments.

### Single-leaf insect choice assay

Single rosette leaves (not detached from the plant) from SAP54 and control plants were fitted opposite each other in a 2cm x 8cm x 12cm (H x W x D) transparent plastic cage fitted with nylon mesh-lined holes (4cm diam.) to allow for air circulation. Five male and 5 female adult *M. quadrilineatus* leafhoppers (which did not carry AY-WB) were introduced into the cage and allowed free access to both leaves. Eggs were dissected and counted under stereomicroscope (15x) five days after the addition of adult insects.

### Statistical analysis

Statistical analysis was performed in Minitab16. Insect oviposition data were analysed using paired t-test, two-tailed t-test or GLM when appropriate. Assumptions of the statistical tests - normal distribution and equal variance - were checked with the Anderson-Darling and the Levene's tests respectively.

## Supplemental information

Supplemental information contains three figures.

**Figure S1.**
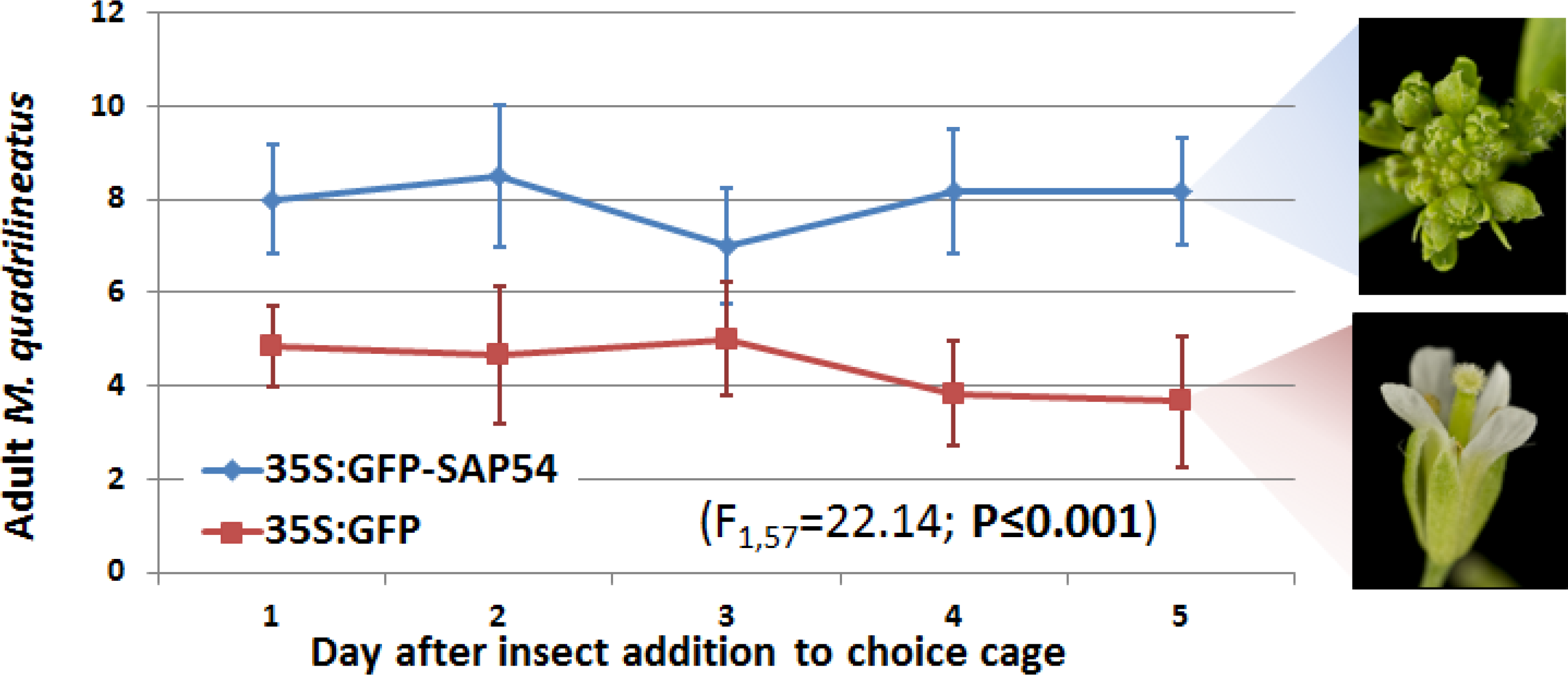
Aster leafhopper *Macrosteles quadrilineatus* demonstrates significantly greater residency preference for SAP54 expressing plants with leaf like-flowers. Plants were grown at 22^o^C, 16/8-hour light/dark to stimulate flowering. Six-week old plants were used for insect choice assay. 10 female and 10 male AY-WB non-infected adult leafhoppers were released in a choice cage containing two tests plants for the period of 5 days. Insects could freely move between the plants. Significantly more insects were found on SAP54 plants over the entire 5 day choice period (GLM with time as covariate; F_157_=22.14; P≤0.001).

**Figure S2.**
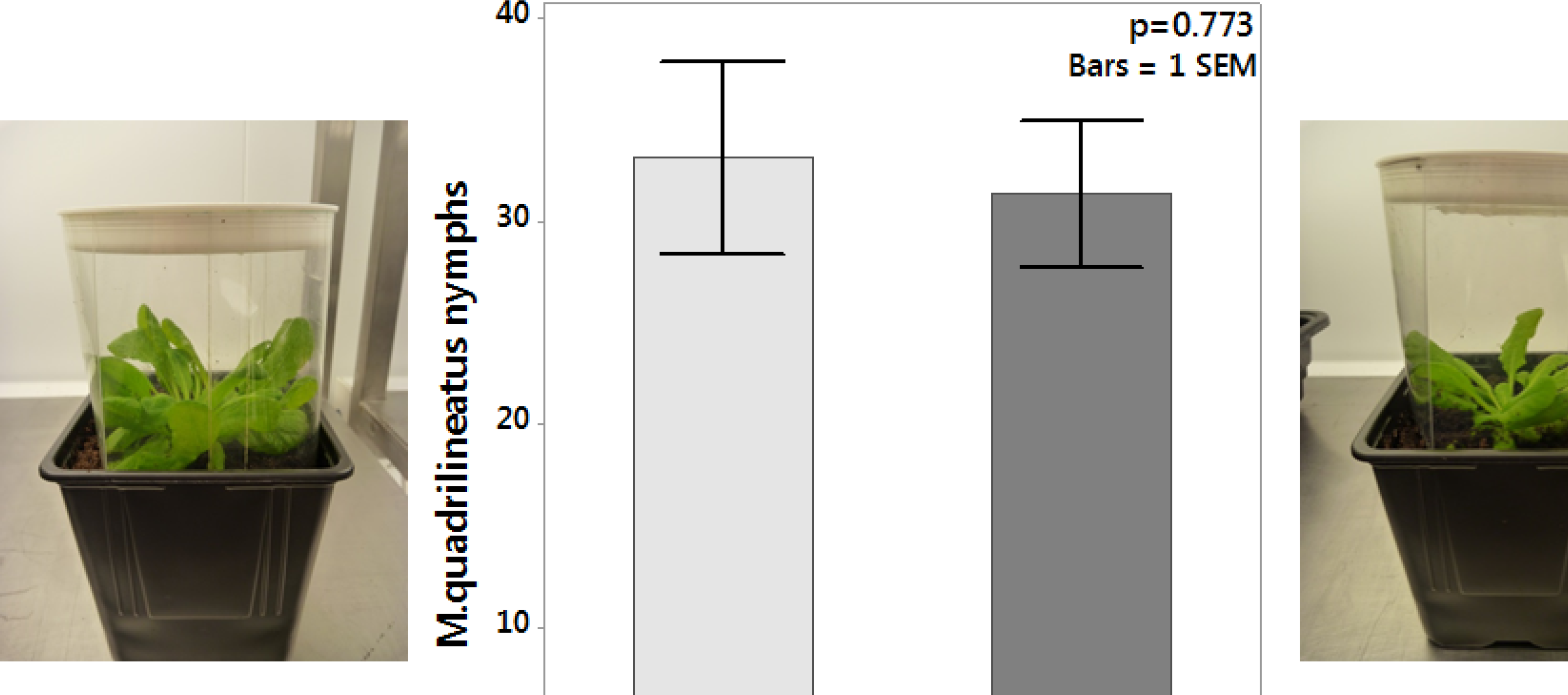
No-choice tests of *M. quadrilineatus* leafhoppers on SAP54 transgenic plants. Equal number of adult *M. quadrilineatus* (5 female + 5 males) were forced to feed and reproduce on SAP54 and control plants to mimic equal host plant selection by insects. Insects were removed 5 days later and nymphs were counted on each test plant 14 days after removal of adult insects from the cages. In contrast to the choice experiments, there was no difference in the number of leafhopper nymphs produced on both plants (paired t-test; n=6; p=0.773).

**Figure S3.**
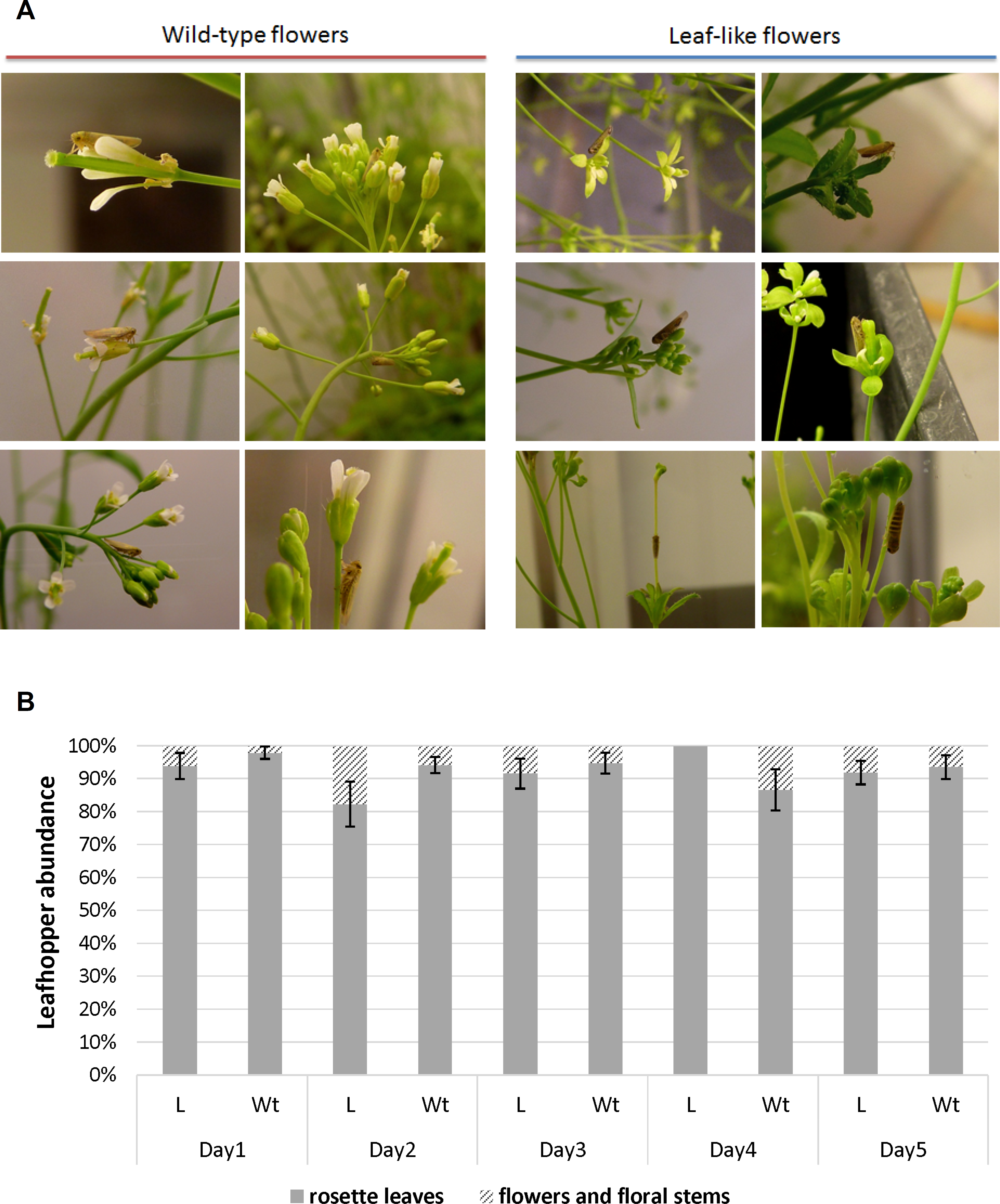
Leafhoppers demonstrate similar distribution on plants with leaflike and wild-type flowers. **A.** *M. quadrilineatus* leafhoppers were photographed whilst residing and feeding on all parts of *Arabidopsis thaliana*(Col-0) plants, including rosette leaves and petioles, stems, cauline leaves and flowers. Insects were found feeding on carpels, sepals, petals and pedicels of wild-type *Arabidopsis thaliana*(Col-0) flowers as well as the leaf-like flowers produced by AY-WB infection or overexpression of a SAP54 alone **B**. A plant with wild type and a plant with leaflike flowers were put in a transparent plastic cage and exposed to 10 male and 10 female adult leafhoppers. Insects were counted separately on floral parts and rosettes for each test plant in the cage once a day. The proportion of insects found on each tissue type is plotted as the mean of 8 replicate cages (bars = 1 SEM). *M. quadrilineatus* has significant residency preference for rosette leaves compared to other floral structures both on AY-WB infected *rad23BCD* mutant plants with leaf-like (L) flowers and AY-WB infected *rad23BD* mutant plants with wild-type flowers (GLM with time as covariate; F_1137_ = 1797.78; P ≤ 0.001). There is no difference between insect residency on wild-type and leaf-like flowers during the five-day leafhopper choice experiment (GLM with time as covariate; F_167_ = 0.19; P = 0.666).

## Acknowledgment

We thank Enrico Coen and Sophien Kamoun for critical feedback on manuscript versions, Richard D. Vierstra (Department of Genetics, University of Wisconsin-Madison) and Martin M. Kater (Department of Bioscience, Universita degli Studi di Milano) for kindly providing seed of *A. thaliana* rad23 mutants and 35S:SVP overexpression lines, respectively. We are grateful to Ian Bedford, Anna Jordan, and Gavin Hatt (JIC Insectary) for rearing and care of leafhopper and phytoplasma stocks and the John Innes Horticultural Services for growing the plants used in this study. We also thank Andrew Davis for plant photographs. The work was funded by BBSRC (BB/J0045531/1) and a BBSRC student fellowship awarded to ZO.

## Authors Contributions

ZO designed the experiments, carried out all experimental work, performed data analysis and drafted the manuscript; SH coordinated the study and helped draft the manuscript. All authors gave final approval for publication.

